# Loss of the endocytic tumor suppressor HD-PTP phenocopies LKB1 and promotes RAS-driven oncogenesis

**DOI:** 10.1101/2023.01.26.525772

**Authors:** Chang-Soo Seong, Chunzi Huang, Austin C. Boese, Yuning Hou, Junghui Koo, Janna K. Mouw, Manali Rupji, Greg Joseph, H. Richard Johnston, Henry Claussen, Jeffrey M. Switchenko, Madhusmita Behera, Michelle Churchman, Jill M. Kolesar, Susanne M. Arnold, Katie Kerrigan, Wallace Akerley, Howard Colman, Margaret A. Johns, Cletus Arciero, Wei Zhou, Adam I. Marcus, Suresh S. Ramalingam, Haian Fu, Melissa Gilbert-Ross

## Abstract

Oncogenic RAS mutations drive aggressive cancers that are difficult to treat in the clinic, and while direct inhibition of the most common KRAS variant in lung adenocarcinoma (G12C) is undergoing clinical evaluation, a wide spectrum of oncogenic RAS variants together make up a large percentage of untargetable lung and GI cancers. Here we report that loss-of-function alterations (mutations and deep deletions) in the gene that encodes HD-PTP (*PTPN23*) occur in up to 14% of lung cancers in the ORIEN Avatar lung cancer cohort, associate with adenosquamous histology, and occur alongside an altered spectrum of KRAS alleles. Furthermore, we show that in publicly available early-stage NSCLC studies loss of HD-PTP is mutually exclusive with loss of LKB1, which suggests they restrict a common oncogenic pathway in early lung tumorigenesis. In support of this, knockdown of HD-PTP in RAS-transformed lung cancer cells is sufficient to promote FAK-dependent invasion. Lastly, knockdown of the Drosophila homolog of HD-PTP (dHD-PTP/Myopic) synergizes to promote RAS-dependent neoplastic progression. Our findings highlight a novel tumor suppressor that can restrict RAS-driven lung cancer oncogenesis and identify a targetable pathway for personalized therapeutic approaches for adenosquamous lung cancer.

## Introduction

Heterogeneous tumor suppressor landscapes are a major driver of therapeutic resistance in RAS-driven cancers^1,2^. In lung cancer, intradriver heterogeneity in KRAS-mutant lung adenocarcinoma (LUAD) is mediated in large part by mutations in co-occurring tumor suppressor genes such as LKB1 (gene *STK11*) and *TP53*. In early-stage RAS-driven lung adenocarcinoma, mutations in tumor suppressors are largely mutually exclusive and thus provide an opportunity to further refine personalized therapeutic approaches.

HD-PTP (gene *PTPN23*) is a highly conserved alternative component of the endosomal sorting required for transport (ESCRT) pathway that acts to sort internalized signaling receptors to multivesicular bodies ^3^. Integrin β1 is a target of HD-PTP and knockdown of HD-PTP in vitro elicits increased FAK activation and cell migration phenotypes. Although HD-PTP has been implicated as a haploinsufficient tumor suppressor in mice, studies from Drosophila show that loss of protein function via mutation leads to hyperplastic overgrowth phenotypes ^4–6^. In this report we used the ORIEN Avatar thoracic cancer cohort to identify an alteration frequency in HD-PTP in up to 14% of lung cancers. Using statistical analyses, we show that mutations and deep deletions (but not shallow deletions) associate with lung adenosquamous histology. Moreover, the frequency of KRAS alleles that normally occur in lung cancer is altered in HD-PTP mutant patients.

In previous work we and others have shown that loss of LKB1 restricts FAK-controlled adhesion signaling, is sufficient to accelerate KRAS-driven tumorigenesis, and is associated with lung adenosquamous histology ^7–10^. Our analysis has revealed that alterations in HD-PTP and LKB1 are mutually exclusive in early-stage LUAD, and more broadly in a large set of solid tumor studies. Furthermore, our studies show that loss of HD-PTP is sufficient to promote FAK-dependent invasion and accelerate tumor progression in combination with a variety of oncogenic RAS alleles in vitro and in vivo.

## Results

### Alterations in the gene that encodes HD-PTP (*PTPN23*) occur in up to 14% of the ORIEN Avatar thoracic cancer cohort

The gene that encodes *HD-PTP, PTPN23*, is found on chromosome 3p21.3 which is frequently hemizygously deleted in lung cancer ^11^. A more recent study has shown that *PTPN23* missense mutations found in public cancer databases destabilize the protein ^5^. However, the consequences and therapeutic implications of *PTPN23* mutations and deep deletions in human cancer remains unexplored. To address this, we analyzed the alteration frequency in *PTPN23* from a collection of publicly available non-small cell lung cancer studies. Interestingly, while the alteration rate for multiple early-stage tumor studies was in the range of 1-2%, a single study of advanced stage lung cancers reported an alteration rate of more than 5% (Fig. 1A). To assess the alteration landscape of *PTPN23* in a larger cohort of more advanced stage lung cancers we used the Oncology Research Information Exchange Network (ORIEN) Avatar database to assess alteration rates in the thoracic (THO) lung cancer cohort. In the ORIEN lung cancer cohort we discovered a *PTPN23* alteration rate as high as 14% depending on histologic subtype (Fig. 1B), with the highest mutation rates found in mucinous and adenosquamous carcinoma. The most identified oncogenic driver mutations were from the RAS pathway with KRAS being mutated in 31% of HD-PTP mutant patients (Fig. 1C). An analysis of KRAS alleles revealed the majority being G12C (38%), followed by G12V (35%), with the remaining split between rare oncogenic KRAS mutations (Fig. 1D).

**Figure 1.**
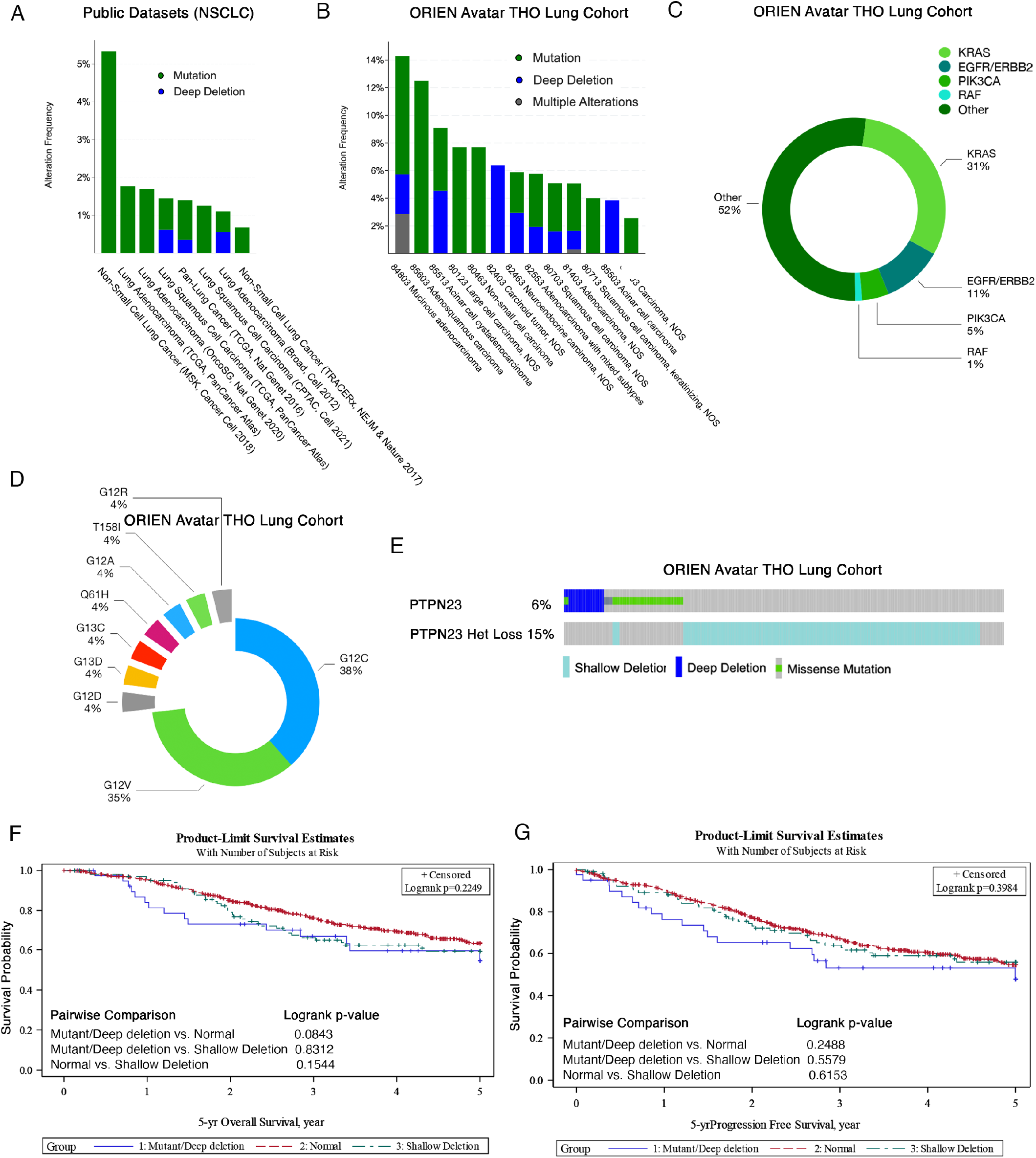
Alteration frequency of HD-PTP in public NSCLC vs. ORIEN Lung Cancer Avatar database. A) Mutation and deep deletion frequency of HD-PTP (gene *PTPN23)* in NSCLC studies from publicly available datasets using cBioPortal ^34,35^ (N = 3,101; stage I/II except for the MSK, 2018 cohort which are advanced stage). B) Mutation and deep deletion frequency of HD-PTP (gene *PTPN23*) in the ORIEN Avatar Thoracic (THO) cancer dataset (N = 1450; stage II-IV). C) Frequency of common receptor tyrosine kinase (RTK) pathway driver mutations in HD-PTP mutant ORIEN THO patients. D) KRAS allele frequency in HD-PTP mutant ORIEN Avatar THO patients. E) Mutual exclusivity analysis of patients with mutations and deep deletions vs. shallow deletions (Het Loss) in the ORIEN Avatar THO patients. (N=862; Log2 Odds Ratio = <-3, p-Value = 0.004 q-Value = 0.004). F) Kaplan-Meier 5 year overall and 5 year progression-free (G) survival estimates for ORIEN Avatar THO patients with different genetic doses of HD-PTP (Group 1: Mutations/Deep deletions; N = 40, Group 2: WT (Normal); N = 597, and Group 3: Shallow Deletions; N = 104). For B-E, data in the January 2023 ORIEN private cBioPortal instance was analyzed.

### Mutations and deep deletions in HD-PTP associate with lung adenosquamous histology

HD-PTP is found on chromosomal arm 3p, which is lost by shallow deletion in a high percentage of cancers, particularly those in the lung ^12^. In addition, haploinsufficiency for HD-PTP predisposes mice to lung cancer and B-cell lymphoma ^5^. In the ORIEN database we found that loss of HD-PTP by shallow deletion is largely mutually exclusive with mutations and deep deletions in HD-PTP (Fig. 1E)). This allowed us to compare clinicopathological associations among HD-PTP wild-type patients, HD-PTP patients with loss of function alterations, and HD-PTP haploinsufficient patients. Data from 753 unique ORIEN Lung Cancer Avatar patients were separated into three groups: 1) WT (no alteration in HD-PTP; n=609), 2) M/D (Mutant/Deep deletion; n=40), and 3) SD (Shallow Deletion; n-=109). Most patients were stage I-II (69.5%), but a significant portion were stage III-IV (30.5%) (Table 1). Major histologic subtypes were adenocarcinoma (48.5%), squamous cell carcinoma (21.6%), and adenosquamous carcinoma (4%). 50.6 % of patients were male and 49.4% were female with the median age at diagnosis being 64.5 [range (19.18-90]. Overall and progression-free survival was estimated between groups. Although the estimates did not reach statistical significance, patients with mutations and deep deletions showed a strong trend in worse overall survival for the pairwise comparison to WT ‘Normal’ (5 year survival rates for M/D = 54.6% [95% CI (35.3%-70.4%)], SD = 59.4% [95% CI (48.1%-69.0%)], WT = 63.5% [95% CI (58.6%-67.9%)]) and a decreased progression free survival, although it did not reach statistical significance (M/D = 47.9% [95% CI (29.5%-64.1%), SD = 55.9% [95% CI (44.7%-65.6%)], WT = 54.7% [95% CI (49.8%-59.3%)) and (Fig. 1F,G).

**Table 1.** Descriptive statistics of ORIEN Avatar THO cancer patients.

| Variable | Level | N (%) = 753 |
| --- | --- | --- |
| Group | Normal | 609 (80.9) |
|  | Mutant/Deep deletion | 40 (5.3) |
|  | Shallow Deletion | 104 (13.8) |
| Smoking status | Never | 91 (13.6) |
|  | Former/ever | 431 (64.3) |
|  | Current | 148 (22.1) |
|  | Missing | 83 |
| Sex | Female | 372 (49.4) |
|  | Male | 381 (50.6) |
| Race | White | 687 (91.5) |
|  | Black | 46 (6.1) |
|  | Others | 18 (2.4) |
|  | Missing | 2 |
| Ethnicity | Spanish, NOS; Hispanic, NOS; Latino, NOS | 15 (2.0) |
|  | Non-Spanish; Non-Hispanic | 726 (98.0) |
|  | Missing | 12 |
| Histology at first Lung CA diagnosis | Adenocarcinoma | 365 (48.5) |
|  | Squamous cell carcinoma | 163 (21.6) |
|  | Adenosquamous carcinoma | 30 (4.0) |
|  | Adenocarcinoma, metastatic, NOS | 1 (0.1) |
|  | Others | 194 (25.8) |
| Clinical Stage at first Lung CA diagnosis | Stage I-II | 455 (69.5) |
|  | Stage III-IV | 200 (30.5) |
|  | Missing | 98 |
| Age at Diagnosis | Mean | 64.31 |
|  | Median | 64.49 |
|  | Minimum | 19.18 |
|  | Maximum | 90.00 |
|  | Std Dev | 10.22 |
|  | Missing | 12.00 |

Clinical covariate analysis between groups indicated a significant difference (P-value <.001) between the major histologic subtypes (Table 2). Mutations/deep deletions (M/D) in HD-PTP were associated with adenosquamous cancers (15% M/D vs. 3.45% WT and 2.88% SD), and shallow deletions in HD-PTP (SD) were associated with squamous morphology (42.31% SD vs. 18.23% WT and 20% M/D). There were no additional associations found between groups for the remaining variables tested. Interestingly, we combined rare architectural subtypes into a single group (Other) and found mutations/deep deletions in HD-PTP were less represented in this group (10% M/D vs. 26.44% WT vs. 27.88% SD).

**Table 2.** Univariate association group with clinicopathological covariates.

| Covariate | Statistics | Level | Group |  |  | P-value |
| --- | --- | --- | --- | --- | --- | --- |
|  |  |  | Normal<br>N=609 | Mutant/Deep<br>deletion N=40 | Shallow<br>Deletion<br>N=104 |  |
| Histology at first<br>Lung CA<br>diagnosis | N (Col %) | Adenocarcinoma | 315 (51.72) | 22 (55) | 28 (26.92) | <.001 |
|  | N (Col %) | Squamous cell<br>carcinoma | 111 (18.23) | 8 (20) | 44 (42.31) |  |
|  | N (Col %) | Adenosquamous<br>carcinoma | 21 (3.45) | 6 (15) | 3 (2.88) |  |
|  | N (Col %) | Adenocarcinoma,<br>metastatic, NOS | 1 (0.16) | 0 (0) | 0 (0) |  |
|  | N (Col %) | Others | 161 (26.44) | 4 (10) | 29 (27.88) |  |
| Smoking status | N (Col %) | Never | 71 (13.25) | 5 (13.51) | 15 (15.46) | 0.953 |
|  | N (Col %) | Former/ever | 344 (64.18) | 24 (64.86) | 63 (64.95) |  |
|  | N (Col %) | Current | 121 (22.57) | 8 (21.62) | 19 (19.59) |  |
| Alcohol use | N (Col %) | Never | 215 (43.43) | 15 (48.39) | 35 (41.67) | 0.202 |
|  | N (Col %) | Former/ever | 41 (8.28) | 6 (19.35) | 9 (10.71) |  |
|  | N (Col %) | Current | 239 (48.28) | 10 (32.26) | 40 (47.62) |  |
| Chronic<br>obstructive<br>pulmonary<br>disease (COPD)<br>diagnosis | N (Col %) | No | 380 (62.4) | 20 (50) | 61 (58.65) | 0.251 |
|  | N (Col %) | Yes | 229 (37.6) | 20 (50) | 43 (41.35) |  |
| Sex | N (Col %) | Female | 300 (49.26) | 22 (55) | 50 (48.08) | 0.749 |
|  | N (Col %) | Male | 309 (50.74) | 18 (45) | 54 (51.92) |  |
| Age at Diagnosis | Median |  | 64.63 | 63.35 | 64.23 | 0.992 |
|  | Min |  | 19.18 | 43.56 | 33.39 |  |
|  | Max |  | 90 | 78.4 | 88.39 |  |

### Loss of HD-PTP leads to hyperactivation of FAK-dependent cell invasion in RAS-mutant lung cancer cell lines

Previous studies have linked loss of HD-PTP to increased cell migration and invasion via Integrin-linked FAK/Src signaling ^6^, but the consequences of loss of HD-PTP in RAS-driven cancers is unknown. To test the functional effects of HD-PTP loss on FAK signaling and cell invasion in human RAS-mutant lung cancer cells, we used both transient and stable knockdown strategies in two HD-PTP wild-type lung cancer cell lines with different RAS drivers (SW1573 (KRAS^G12C)^ and H1299 (NRAS^G61L^). An average knockdown efficiency of 94.5% was achieved across all experiments. Knockdown of HD-PTP was sufficient to upregulate Integrin β1 levels and FAK activity in both cell lines (Fig. 2A). To test whether upregulation of FAK resulted in functional changes in cell behavior we tested the effect of HD-PTP knockdown on cell invasion in 2D. Knockdown of HD-PTP resulted in significant changes in 2D invasion, with no major effects on cell growth (Fig. 2B-D). Since cancer cells grown in 3D culture conditions more faithfully mimic critical aspects of tumor architecture important in predicting drug response ^13^ we tested whether loss of HD-PTP promotes 3D invasion of RAS-mutant lung cancer spheroids. Loss of HD-PTP in both SW1573 and H1299 cells increased invasive area and decreased circularity (and indirect measure of sheet-like collective invasion ^14^ in each cell line using both transient and stable knockdown methods (Fig. 2E-H). In prior published work, we have shown that FAK-dependent collective invasion in vivo and in 3D cell culture can be suppressed using a pharmacologic FAK inhibitor ^9^. To test if FAK inhibition can suppress 3D invasion in RAS/HD-PTP-mutant tumor cells we treated invading spheroids with the FAK inhibitor defactinib. Treatment of 3D spheroids with defactinib was sufficient to suppress FAK activity and inhibit increased collective invasion associated with HD-PTP knockdown in both H1299 and SW1573 cells (Fig. 2I,J).

**Figure 2.**
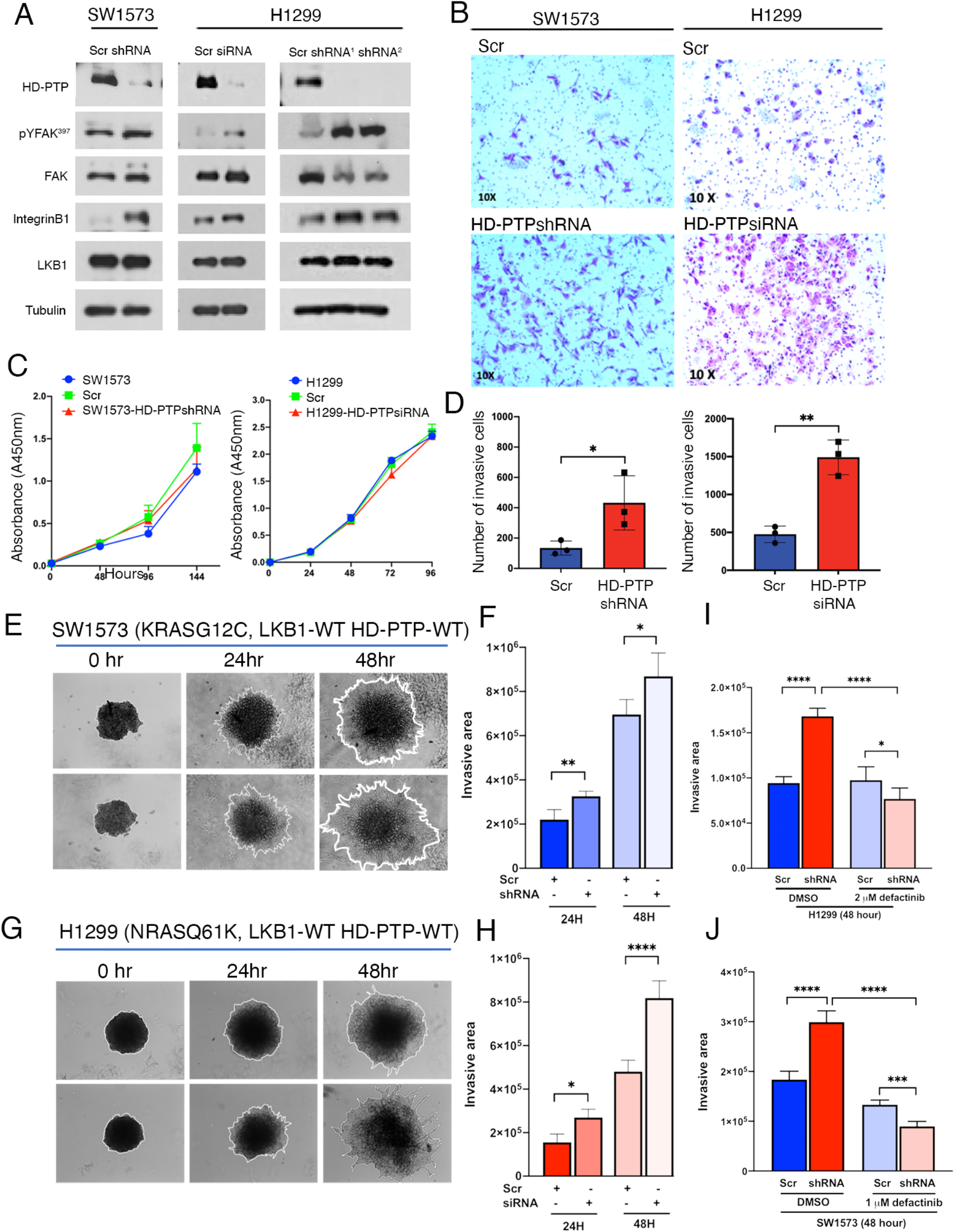
Knockdown of HD-PTP is sufficient to activate FAK-dependent invasion in RAS-mutant lung cancer cell lines. A) Western blot of indicated signaling proteins in SW1573 (KRAS^G12C^/LKB1^WT^) and H1299 (NRAS^Q61K^/LKB1^WT^) lung cancer cells expressing either scrambled control, HD-PTPshRNA or HD-PTP siRNA. B) Brightfield images of 2D invasion assays of SW1573 and H1299 cells expressing scrambled control and shRNA or siRNA to HD-PTP. C) Growth of cell populations expressing either shRNA or siRNA to HD-PTP. D) Quantitation of invading cells from 2D invasion assays in (B). 2D invasion assays were performed using three biological replicates. Error bars represent the Std. Deviation of Mean. Statistical significance was analyzed using the student’s t-test. * = <.05; ** = <.01. E) Representative images of SW1573 3D spheroids expressing either scrambled control or HD-PTP shRNA and embedded in invasion matrix for the indicated time. Quantification of invasive area in the SW1573 assay in (E). G) Representative images of H1299 spheroids expressing scrambled control or siRNA to HD-PTP and embedded in invasion matrix for the indicated time. H) Quantification of invasive area in the H1299 invasion assay in (G). Quantification of invasive area in H1299 (I) and SW1573 (J) cells expressing scrambled control or shRNA to HD-PTP and treated with vehicle control (DMSO) or the FAK inhibitor defactinib. Quantitative data were analyzed using 4-5 biological replicates. Error bars represent Std. Deviation of Mean and statistical significance was tested using one-way ANOVA, multiple comparisons. * = <.05; ** = <.01; **** = <.0001.

### HD-PTP mutations are mutually exclusive of LKB1 in early-stage NSCLC and deregulate integrin-linked adhesion signaling in vivo

Mutations in the master kinase and tumor suppressor LKB1 co-segregate with oncogenic KRAS, restrict FAK-controlled adhesion signaling, and are sufficient to promote the progression of latent Kras lung tumor in vivo ^7–10,15^. The shared phenotypes between loss of HD-PTP and LKB1 in lung cancer cells prompted us to hypothesize that mutations would be mutually exclusive of one another. By analyzing early-stage non-small cell lung cancers from publicly available studies we discovered that alterations in the gene that encodes LKB1 (*STK11*) are mutually exclusive with alterations in the gene that encodes HD-PTP (*PTPN23*), both in a smaller set of non-small cell lung cancer studies, and more broadly in a large set of non-redundant solid tumor studies (Fig. 1A,B). These data suggest HD-PTP and LKB1 regulate a common signaling pathway that is critical to tumorigenesis.

In previous studies, we isolated loss-of-function alleles of the Drosophila homolog of HD-PTP (dHD-PTP/Myopic) in a screen for conditional growth suppressors ^4^. If HD-PTP and LKB1 do indeed regulate a common signaling pathway to restrict tumorigenesis, we would predict that loss of HD-PTP, like LKB1, would synergize with oncogenic RAS to promote tumor progression in vivo. To begin to test this using an in vivo Drosophila model, we first verified that alterations in Drosophila HD-PTP lead to integrin-linked adhesion phenotypes. Since homozygous mutant dHD-PTP epithelial clones in imaginal discs undergo apoptosis, we used the H99 deletion to block cell death. In control *H99* clones the Drosophila homolog of Integrin ß1, ßPS integrin, is concentrated at the basal side of epithelial cells (Fig. 3C). In H99/dHD-PTP mutant clones, ßPS integrin mislocalizes and accumulates to the apical surface, which phenocopies loss of HD-PTP in mammalian cells ^16^. Moreover, overexpression of dHD-PTP in the posterior compartment of the Drosophila wing phenocopied overexpression and activation of Drosophila FAK (Fak56D) and constitutively active ßPS integrin by resulting in cell adhesion defects visible as wing ‘blisters’ (Fig. 3D). These data suggest the ability of HD-PTP to limit integrin/FAK signaling through sorting activated integrin complexes is conserved from Drosophila to humans.

**Figure 3.**
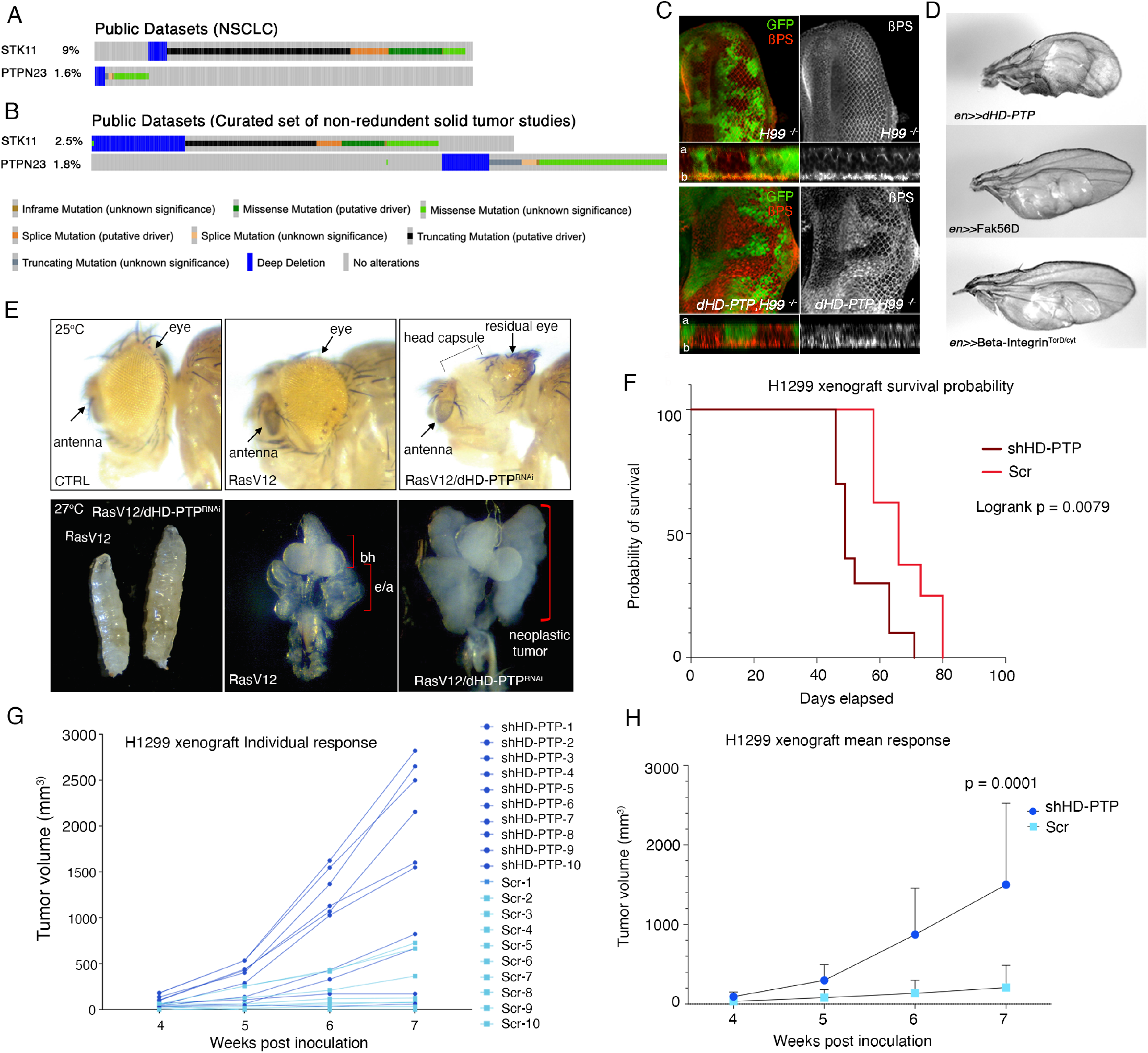
Mutations in the gene that encodes HD-PTP are mutually exclusive with those of LKB1 and synergize in vivo to promote RAS-driven tumor progression. (A) Oncoprint generated from public cBioPortal data representing mutual exclusivity analysis of 3,101 NSCLC cases the majority of which are lung adenocarcinoma ^36–40^; N = 3101, Log2 Odds Ratio == <-3; p-value = 0.011. (B) Oncoprint generated from public cBioPortal representing mutual exclusivity analysis early-stage non-redundant solid tumor studies {CITE}; N = 23,688, Log2 Odds Ratio = -2.285; p-value = 0.047; q-value = 0.047). C) Representative confocal images of Drosophila 3^rd^ instar larval eye imaginal discs carrying control H99 clones (marked by the absence of GFP) (top) or discs carrying *dHD-PTP,H99* clones (marked by the absence of GFP) and stained for βPS integrin. a = apical; b = basolateral. D) Representative images of adult Drosophila wings over-expressing the indicated transgenes in the posterior wing compartment (*en>Gal4*). E) Brightfield images of adult Drosophila eyes expressing low levels (25°) of the indicated transgenes (top). High-level expression of RasV12/dHD-PTP^RNAi^ (27°) leads to a ‘giant’ larva phenotype and neoplastic tumors that kill the animal (bottom row). e/a = eye/antennal; bh= brain hemispheres. F) Kaplan-meier analysis of H1299 mouse xenografts expressing either Scr control (8 mice/group; mean survival = 66 days) or shHD-PTP (10 mice/group; mean survival = 49 days). Individual growth responses from H1299 mouse xenografts expressing either Scr control (10 mice/group) or shHD-PTP (10 mice/group). H) Mean growth response from H1299 mouse xenografts expressing either Scr control (10 mice/group) or shHD-PTP (10 mice/group). Survival estimates were analyzed for statistical significance using Log-rank test, and mean tumor volumes were analyzed using 2way ANOVA with Sidak’s multiple comparisons test.

### Loss of HD-PTP is sufficient to promote the progression of latent RAS-driven tumors in vivo

We recently reported that tumorigenic cooperation between oncogenic Ras and loss of Lkb1 is conserved in Drosophila ^17^. To test whether loss of dHD-PTP is sufficient to promote the neoplastic progression of tissues expressing oncogenic Ras, we used the temperature sensitive Gal4 system in early eye progenitor cells to express both low and high levels of oncogenic Ras (RasV12) and a validated dHD-PTP/Mop RNAi transgene. Low level expression of oncogenic Ras resulted in a slightly hyperplastic eye (Fig. 3E) and knockdown of dHD-PTP in these cells results in a stronger phenotype of transdifferentiating eye tissue and benign cuticular overgrowth. Conversely, compared to high-level expression of RasV12 alone, knockdown of dHD-PTP and high-level RasV12 results in a ‘giant’ larval phenotype with grossly overgrown neoplastic eye/imaginal disc tumors that kill the animal. To validate our results in a mammalian system we established H1299 lung cancer xenograft tumors expressing either scrambled control or HD-PTPshRNA. Knockdown of HD-PTP in vivo resulted accelerated tumor growth and decreased survival (Fig. 3F-H). These data suggest loss of HD-PTP can cooperate with oncogenic RAS to promote tumorigenesis in vivo.

## Discussion

RAS mutations occur in aggressive tumors and confer a high variability in therapeutic response. Co-occurring tumor suppressor mutations like LKB1 and TP53 are major effectors of subtype-specific biology and outcomes. In this report, we show that loss of HD-PTP phenocopies LKB1 by restricting adhesion signaling and invasion in RAS-transformed lung cells, and by promoting tumor progression in a RasV12-driven Drosophila model. Our analysis of the ORIEN Avatar THO lung cancer cohort shows that HD-PTP (gene *PTPN23*) is altered in a high percentage of stage II-IV lung cancers (up to 14%) and associates with adenosquamous morphology. Moreover, in stage I-II NSCLC patients and a large cohort of non-redundant solid tumor studies alterations in HD-PTP are mutually exclusive with those of LKB1, further supporting our conclusion that HD-PTP and LKB1 regulate a common pathway in early oncogenesis.

At its N-terminus, HD-PTP contains a well-conserved Bro domain that is required, in conjunction with the endosomal sorting complex required for transport (ESCRT) complex, for the sorting of internalized cell surface receptors to multivesicular bodies (MVBs) and is thus functionally homologous to dHD-PTP (Myopic) ^6^. Multiple lines of evidence exist that the PTP domain at the C-terminus of HD-PTP is a pseudophosphatase domain. Although the cysteine required for enzymatic activity is conserved, the invariant alanine normally found at position +2 in the PTP active site has been replaced by a serine and the Asp residue that acts as a general acid has been replaced by Glu. Moreover, functional experiments have shown that HD-PTP does not exhibit detectable phosphatase activity against a panel of sensitive protein and lipid substrates ^18,19^. This suggests HD-PTP/Myopic restricts FAK activity through its endolysosomal sorting function of activated integrin receptors, which is supported by our data showing upregulated βPS integrin on the apical surface of mop clones and by prior studies of human HD-PTP ^16,20^.

We show that mutations and deep deletions in HD-PTP associate with adenosquamous lung histology, a phenotype shared by LKB1. Notably, knockdown of dHD-PTP in low-level RasV12-expressing eye progenitor cells results in transdifferentiation into surrounding head capsule and cuticle structures. HD-PTP is a conserved regulator of multiple developmental signaling pathways. In our previous published work we identified Yorkie, the Drosophila homolog of the Hippo pathway transcriptional co-activator, as a physical binding partner and mediator of dHD-PTP-mutant overgrowth phenotypes ^4^. Mammalian YAP is an oncogene and cell fate determinate and has been shown to regulate malignant progression and cell lineage transition of Lkb1-deficient LUAD ^21–24^. Future studies should focus on whether loss of HD-PTP is sufficient to alter tumor cell fate, and identifying the cell-fate determinate regulated by HD-PTP in lung cells.

Lastly, our show that loss of HD-PTP in RAS-transformed lung cancer cells leads to upregulated Integrin-linked FAK activation and increased cell invasion. Hyperactive FAK is associated with poor survival, increased invasion and metastasis, and has been implicated in the resistance to immune checkpoint therapy ^25,26^. Our previous work has shown that KRAS/LKB1-mutant lung cancers respond to FAK inhibition in vivo ^9^. FAK inhibitors have shown promise in novel combination regimens ^27^. Therefore, combination treatment strategies involving FAK pathway inhibition may be a viable treatment option for HD-PTP mutant patients, including those with rare and treatment resistant adenosquamous histology.

### Limitations of the study

Loss of HD-PTP and LKB1 are not mutually exclusive in the ORIEN THO lung cancer Avatars. We hypothesize this is due to the larger number of late and advanced stage patients in this cohort. Moreover, mutual exclusivity became apparent in the large set of non-redundant solid tumor studies after removing those solely focused on metastatic or treatment refractory tumors. We did not detect mutual exclusivity between loss of HD-PTP and P53 even in early-stage cohorts. Therefore, the biological function shared by LKB1 and P53 in early tumorigenesis is unlikely shared by HD-PTP.

## Methods

### ORIEN Avatar THO lung cancer cohort

ORIEN is a cancer precision medicine initiative initially developed by the Moffitt Cancer Center ^28,29^. It has evolved into a consortium research network of nineteen U.S. cancer centers. All ORIEN alliance members utilize a standard protocol: Total Cancer Care (TCC)^®^. TCC is a prospective cohort study with whole-exome tumor sequencing, RNA sequencing, germline sequencing, and lifetime follow up. Nationally, over 250,000 participants have enrolled.

### Mutation and copy number analysis

792 ORIEN samples with RNAseq fastq files were returned to the PI. Samples were bucketed based on mutational status of the PTPN23 gene. ORIEN analysis had previously identified 33 individuals with a SNP or small INDEL in the PTPN23 gene. CNV analysis was performed by ORIEN, using CNTools, to identify samples with 0 copies of PTPN23 (deep deletion), 1 copy of PTPN23 (shallow deletion), or 2 copies of PTPN23 (no mutation). All the SNP/small INDEL samples had 2 copies of PTPN23, so there was no overlap in the groups. In the end, we analyzed 609 patients with no PTPN23 mutation, 40 with a SNP/small INDEL or deep deletion, and 104 with a shallow deletion.

### Statistics and Analysis

Descriptive statistics were generated for all patient characteristics. Frequency and percentage were reported for categorical variables, and mean, median, standard deviation, IQR, and range were reported for numeric variables. Differences between the study groups for each clinicopathological variable was assessed using chi-sq. test or fisher’s exact test for categorical variables and ANOVA for numeric variables. 5-year Progression Free Survival (PFS) was defined from diagnosis by calculating the difference between age at diagnosis and age at first progression or death which ever came earlier for those with an event or age at last contact for those censored. PFS estimated using the Kaplan-Meier method, and PFS was compared using log-rank tests. Univariate (UVA) cox regression analysis was used to determine the effect of each clinicopathological variable with PFS. A multivariable cox regression analysis by a backward selection method was used to select the covariates applying an alpha of 0.05. 5-year overall survival was defined from diagnosis and estimated as the difference between age at first lung diagnosis and age at death for those with an event and age at last contact for those censored. MVA were performed like above. Statistical analysis was performed using SAS 9.4 (SAS Institute Inc., Cary, NC) macros ^30^, and statistical significance was assessed at the 0.05 level.

### Western blotting

Cells were lysed in 1X lysis buffer (Cell Signaling) and 30 ug of cell lysates were resolved by 10% SDS-PAGE gel and transferred overnight to a PDVF membrane at 0.07A at 4°C. Membranes were blocked in 5% BSA or 5% milk for one hour at room temperature and then incubated overnight at 4°C with rabbit anti-HD-PTP (1:1,000, Bethyl Laboratories), mouse anti-Integrinβ1/CD29(1:1,000 BD Biosciences), rabbit anti-phospho FAK(Tyr397) (1:1,000, Invitrogen), rabbit-anti-FAK(1:1,000, Cell Signaling) and rabbit anti-LKB1 (1:1000, Cell Signaling) in TBS-T (0.1% Tween 20) containing 5% BSA or 3% milk. Tubulin was detected as a loading control by incubation of the membrane with the E7 antibody from Developmental Studies Hybridoma Bank at a 0.2 ug/ml dilution for overnight at 4°C. Membranes were washed in TBS-T, incubated overnight with primary antibodies at 4 degrees and detected with goat anti-rabbit or mouse-HRP secondary antibodies (Thermo Scientific) followed by detection using SuperSignal™ West Pico Plus chemiluminescent substrate (Thermo Scientific).

### 2D invasion assays

Transwell invasion assays were performed using inserts with 8.0-μm pores (Falcon). 5 × 10^4^ cells were mixed in total 5 mg/ml growth factor reduced Matrigel matrix (Corning) and plated in inserts in a total volume of 30ul, and inserts were placed in 24-well plates (Corning). After incubation for 30 min at 37°C in a 5% CO_2_ incubator, 200 ul serum-free culture medium was added to the inserts followed by the addition of 500 ul of complete culture media to the bottom wells. After 24 hr, invaded cells that accumulated on the bottom surface of the insert were fixed with 4% paraformaldehyde for 10 min at room temperature, washed with 1× phosphate-buffered saline (PBS) for 10 min. The cells were then fixed with 100% methanol for 10 min at room temperature, washed 3 times with 1× PBS and stained with 0.5% Crystal violet for 30 min. The cells and Matrigel were cleaned out of the inserts using cotton swabs and the membranes were mounted on slides and imaged using an Olympus CKX41 inverted microscope and analyzed using Infinity Analyze 7. The total number of cells/membrane were counted and the experiments were performed each in triplicate.

### Cell growth assays

Cell counting kit-8(CCK-8, Dojindo) was used to assay cell growth. 1× 10^3^ cells were grown in 96-well plates for 24hr through 144 hours. Every 24 h term, cell growth was detected using a CCK-8 assay kit followed by company instruct protocol. 10 µl of CCK-8 solution was added to each well, followed by incubation for 3 h at 37°C. The absorbance at 450 nm was determined by a multiplate reader (BioTek). The mean value and STDEV from 4 wells were calculated.

### 3D spheroid invasion assays and drug treatments

Cells were suspended in RPMI with 10% FBS in 96-well round bottom plates at a density of 3× 10^3^ cells/well. The cells in round-bottom plates were centrifuged at 1500 rpm for 10 min and incubated at 37 °C in a 5% CO_2_ incubator for 72hrs to form single spheroids. Spheroids were transferred into 35mm glass bottom dishes and embedded in 5mg/ml growth factor reduced Matrigel with media contained DMSO, 1uM or 2uM defactinib. After 30 minutes of incubation at 37º C with 5% CO_2_ spheroids were overlayed with 2mL of pre-warmed culture media alone, DMSO, 1uM or 2uM defactinib. Images were captured at time 0 and every 24hrs for 3 days using an Olympus CKX41 inverted microscope. Invasive area was calculated by measuring the difference between the total spheroid area and the spheroid core in Image J. Spheroid circularity was utilized as an indirect measure of sheet-like collective invasion and was quantified in Image J.

### Generation of HD-PTP knockdown cell lines

SW1573 cells (ATCC) were grown in L-15(ATCC 30-2008) medium with 10% FBS, 2mM l-glutamine, and 1% penicillin-streptomycin. H1299 cells (ATCC) were grown in RPMI 1640 with 10% FBS, 2mM l-glutamine, and 1% penicillin-streptomycin. For transient knockdown, scrambled non-targeting (D-001210-01-05; siGENOME) and HD-PTP (*PTPN23*) siRNA (D-009417-01-0005; siGENOME; Horizon) were transfected using Lipofectamine RNAiMax reagent (Thermofisher). To generate stable HD-PTP knockdown cell lines we transduced SW1573 and H1299 cells with 3 individual HD-PTP shRNAs (LPP-HSH067569-LVRH1GH) and control lentiviral particles (Scramble, LPP-CSHCTR001-LVRH1GH) (GeneCopoeia). Cells were selected with hygromycin and the strongest HD-PTP shRNA knockdown was used for the remaining experiments. The FAK inhibitor defactinib (VS-6063) was purchased from Selleckchem.

### Drosophila genetics

Flies were raised on a standard cornmeal/agar molasses medium at 25 °C with 50% relative humidity under a 12hr light/12hr dark cycle. The *dHP-PTP* (*mop*^sfv3^),*Df(3L)H99,FRT80B/TM6B, Df(3L)H99,FRT80B/TM6B* ^4^ and *eyFLP; ubi:GFP,FRT80B* (Bloomington Drosophila Stock Center (BDSC) lines were used to generate control and *dHD-PTP* homozygous null clones within an otherwise heterozygous animal. UAS-Mop ^31^, *UAS-Fak56D* ^32^ and *UAS-ß-Integrin*^*TorD/Cyt*33^ and *en-Gal4, UAS-GFP*.*nls/CyO* (BDSC) were used for misexpression studies. dHD-PTP (*myopic* (*mop*)) RNAi line (UAS-Mop^Trip20^; #32916) was obtained from the Bloomington Drosophila Stock Center (BDSC). *ey-GAL4* (#5535) and UAS-Ras^V12^ (#64196) were also provided by BDSC. All images were captured using a Leica X6D microscope, digital camera, and Leica Acquire software.

### Xenograft assays

1 × 10^^6^ stable H1299 Scr control or H1299-HD-PTPshRNA cells were prepared in 25% Matrigel/PBS and injected subcutaneously into NSG mice (10 mice/group). Post-injection, mice were examined regularly for palpable tumors. Once palpable, tumor volume measurements were collected weekly using Vernier calipers until mice reached Emory University IACUC endpoint guidelines.

## Acknowledgements

We would like to thank the following ORIEN Cancer Center Sites for contributing THO-Lung Cancer Avatar cases: H. Lee Moffitt Cancer Center & Research Institute, The Ohio State University Comprehensive Cancer Center, University of Virginia Cancer Center, University of Colorado Cancer Center, Rutgers Cancer Institute of New Jersey, USC Norris Comprehensive Cancer Center, Huntsman Cancer Institute, Oklahoma University Stephenson Cancer Center, Roswell Park Comprehensive Cancer Center, The University of Kentucky Markey Cancer Center, and Indiana University Simon Cancer Center. Research reported in this publication was supported by the National Cancer Institute of the National Institutes of Health under Award Number R01CA194027 (MGR, WZ, AIM), R01CA201340 (MGR, AIM), R01CA250422 (AIM), R01CA236369 (AIM), P50CA217691 (SSR, HF), and Winship Cancer Institute Developmental Funds. Research reported in this publication was supported in part by the Emory Integrated Cellular Imaging Core, the Emory Integrated Genomics Core, and the Cancer Animal Models, Winship Data and Technology Applications, and Biostatistics and Bioinformatics Shared Resource of the Winship Cancer Institute of Emory University and NIH/NCI under award number P30CA138292 (SSR). The content is solely the responsibility of the authors and does not necessarily represent the official views of the National Institutes of Health.

## Author contributions

Chang-Soo Seong: Conceptualization, Methodology, Investigation, Writing - Original Draft, Visualization

Chunzi Huang: Methodology, Investigation

Austin C. Boese: Conceptualization, Formal analysis

Yuning Hou: Methodology, Formal Analysis

Junghui Koo: Methodology, Resources

Janna K. Mouw: Methodology, Resources

Manali Rupji: Formal analysis

Greg Joseph: Software

Jeffrey M. Switchenko: Formal analysis

Madhusmita Behera: Project Administration

Michelle Churchman: Resources, Project Administration

H. Richard Johnston: Methodology, Data Curation, Formal Analysis

Henry Claussen: Methodology, Formal Analysis

Margaret A. Johns: Resources, Data Curation, Project administration

Jill M. Kolesar: Resources, Project Administration

Susanne M. Arnold: Project administration, Resources

Katie Kerrigan: Project administration, Resources

Wallace Akerley: Project administration, Resources

Howard Colman: Project administration, Resources

Wei Zhou: Formal Analysis, Resources, Funding acquisition

Adam I. Marcus: Resources, Funding acquisition

Suresh Ramalingam: Funding acquisition

Haian Fu: Funding acquisition

Melissa Gilbert-Ross: Conceptualization, Methodology, Formal analysis, Investigation, Writing - Original Draft, Visualization, Supervision, Project administration, Funding acquisition.

## References

1. Ryan, M.B., and Corcoran, R.B. (2018). Therapeutic strategies to target RAS-mutant cancers. Nature Reviews Clinical Oncology, 1–12.

2. Désage, A.-L., Léonce, C., Swalduz, A., and Ortiz-Cuaran, S. (2022). Targeting KRAS Mutant in Non-Small Cell Lung Cancer: Novel Insights Into Therapeutic Strategies. Frontiers Oncol 12, 796832. 10.3389/fonc.2022.796832.

3. Desrochers, G., Kazan, J.M., and Pause, A. (2019). Structure and functions of His domain protein tyrosine phosphatase in receptor trafficking and cancer1. Biochem Cell Biol 97, 68–72.

4. Gilbert, M.M., Tipping, M., Veraksa, A., and Moberg, K.H. (2011). A Screen for Conditional Growth Suppressor Genes Identifies the Drosophila Homolog of HD-PTP as a Regulator of the Oncoprotein Yorkie. Developmental cell 20, 700–712.

5. Manteghi, S., Gingras, M.-C., Kharitidi, D., Galarneau, L., Marques, M., Yan, M., Cencic, R., Robert, F., Paquet, M., Witcher, M., et al. (2016). Haploinsufficiency of the ESCRT Component HD-PTP Predisposes to Cancer. Cell reports 15, 1893–1900.

6. Gingras, M.-C., Kazan, J.M., and Pause, A. (2017). Role of ESCRT component HD-PTP/PTPN23in cancer. Biochemical Society transactions 45, 845–854.

7. Carretero, J., Shimamura, T., Rikova, K., Jackson, A.L., Wilkerson, M.D., Borgman, C.L., Buttarazzi, M.S., Sanofsky, B.A., McNamara, K.L., Brandstetter, K.A., et al. (2010). Integrative Genomic and Proteomic Analyses Identify Targets for Lkb1-Deficient Metastatic Lung Tumors. CCELL 17, 547–559.

8. Kline, E.R., Shupe, J., Gilbert-Ross, M., Zhou, W., and Marcus, A.I. (2013). LKB1 represses focal adhesion kinase (FAK) signaling via a FAK-LKB1 complex to regulate FAK site maturation and directional persistence. The Journal of biological chemistry 288, 17663–17674.

9. Gilbert-Ross, M., Konen, J., Koo, J., Shupe, J., Robinson, B.S., Wiles, W.G., Huang, C., Martin, W.D., Behera, M., Smith, G.H., et al. (2017). Targeting adhesion signaling in KRAS, LKB1 mutant lung adenocarcinoma. JCI insight 2, e90487. 10.1172/jci.insight.90487.

10. Skoulidis, F., and Heymach, J.V. (2019). Co-occurring genomic alterations in non-small-cell lung cancer biology and therapy. Nature Reviews Cancer, 1–15.

11. Toyooka, S., Ouchida, M., Jitsumori, Y., Tsukuda, K., Sakai, A., Nakamura, A., Shimizu, N., and Shimizu, K. (2000). HD-PTP: A novel protein tyrosine phosphatase gene on human chromosome 3p21.3. Biochemical and Biophysical Research Communications 278, 671–678.

12. Zabarovsky, E.R., Lerman, M.I., and Minna, J.D. (2002). Tumor suppressor genes on chromosome 3p involved in the pathogenesis of lung and other cancers. Oncogene 21, 6915–6935.

13. Langhans, S.A. (2018). Three-Dimensional in Vitro Cell Culture Models in Drug Discovery and Drug Repositioning. Front Pharmacol 9, 6.

14. Konen, J., Summerbell, E., Dwivedi, B., Galior, K., Hou, Y., Rusnak, L., Chen, A., Saltz, J., Zhou, W., Boise, L.H., et al. (2017). Image-guided genomics of phenotypically heterogeneous populations reveals vascular signalling during symbiotic collective cancer invasion. Nat Commun 8, 15078.

15. Ji, H., Ramsey, M.R., Hayes, D.N., Fan, C., McNamara, K., Kozlowski, P., Torrice, C., Wu, M.C., Shimamura, T., Perera, S.A., et al. (2007). LKB1 modulates lung cancer differentiation and metastasis. Nature 448, 807–810.

16. Kharitidi, D., Apaja, P.M., Manteghi, S., Suzuki, K., Malitskaya, E., Roldan, A., Gingras, M.-C., Takagi, J., Lukacs, G.L., and Pause, A. (2015). Interplay of Endosomal pH and Ligand Occupancy in Integrin α5β1 Ubiquitination, Endocytic Sorting, and Cell Migration. Cell Reports 13, 599–609.

17. Rackley, B., Seong, C.-S., Kiely, E., Parker, R.E., Rupji, M., Dwivedi, B., Heddleston, J.M., Giang, W., Anthony, N., Chew, T.-L., et al. (2021). The level of oncogenic Ras determines the malignant transformation of Lkb1 mutant tissue in vivo. Commun Biology 4, 142.

18. Gingras, M.-C., Zhang, Y.L., Kharitidi, D., Barr, A.J., Knapp, S., Tremblay, M.L., and Pause, A. (2009). HD-PTP Is a Catalytically Inactive Tyrosine Phosphatase Due to a Conserved Divergence in Its Phosphatase Domain. PLoS ONE 4, e5105. 10.1371/journal.pone.0005105.

19. Barr, A.J., Ugochukwu, E., Lee, W.H., King, O.N.F., Filippakopoulos, P., Alfano, I., Savitsky, P., Burgess-Brown, N.A., Müller, S., and Knapp, S. (2009). Large-Scale Structural Analysis of the Classical Human Protein Tyrosine Phosphatome. Cell 136, 352–363.

20. Chen, D.Y., Li, M.Y., Wu, S.Y., Lin, Y.L., Tsai, S.P., Lai, P.L., Lin, Y.T., Kuo, J.C., Meng, T.C., and Chen, G.C. (2012). The Bro1-domain-containing protein Myopic/HDPTP coordinates with Rab4 to regulate cell adhesion and migration. Journal of Cell Science 125, 4841–4852.

21. Nguyen, H.B., Babcock, J.T., Wells, C.D., and Quilliam, L.A. (2012). LKB1 tumor suppressor regulates AMP kinase/mTOR-independent cell growth and proliferation via the phosphorylation of Yap. Oncogene. 10.1038/onc.2012.431.

22. Mohseni, M., Sun, J., Lau, A., Curtis, S., Goldsmith, J., Fox, V.L., Wei, C., Frazier, M., Samson, O., Wong, K.-K., et al. (2014). A genetic screen identifies an LKB1-MARK signalling axis controlling the Hippo-YAP pathway. Nature Cell Biology 16, 108–117.

23. Gao, Y., Zhang, W., Han, X., Li, F., Wang, X., Wang, R., Fang, Z., Tong, X., Yao, S., Li, F., et al. (2014). YAP inhibits squamous transdifferentiation of Lkb1-deficient lung adenocarcinoma through ZEB2-dependent DNp63 repression. Nat Commun 5, 4629.

24. Zhang, W., Gao, Y., Li, F., Tong, X., Ren, Y., Han, X., Yao, S., Long, F., Yang, Z., Fan, H., et al. (2015). YAP Promotes Malignant Progression of Lkb1-Deficient Lung Adenocarcinoma through Downstream Regulation of Survivin. Cancer Res 75, 4450–4457.

25. Sulzmaier, F.J., Jean, C., and Schlaepfer, D.D. (2014). FAK in cancer: mechanistic findings and clinical applications. Nature Reviews Cancer 14, 598–610.

26. Jiang, H., Hegde, S., Knolhoff, B.L., Zhu, Y., Herndon, J.M., Meyer, M.A., Nywening, T.M., Hawkins, W.G., Shapiro, I.M., Weaver, D.T., et al. (2016). Targeting focal adhesion kinase renders pancreatic cancers responsive to checkpoint immunotherapy. Nature Medicine 22, 851–860.

27. Dawson, J.C., Serrels, A., Stupack, D.G., Schlaepfer, D.D., and Frame, M.C. (2021). Targeting FAK in anticancer combination therapies. Nat Rev Cancer 21, 313–324.

28. Fenstermacher, D.A., Wenham, R.M., Rollison, D.E., and Dalton, W.S. (2011). Implementing Personalized Medicine in a Cancer Center. Cancer J 17, 528–536.

29. Caligiuri, M.A., Dalton, W.S., Rodriguez, L., Sellers, T., and Willman, C.L. (2016). Orien: Reshaping Cancer Research & Treatment. Oncol Issues 31, 62–66.

30. Liu, Y., Nickleach, D.C., Zhang, C., Switchenko, J.M., and Kowalski, J. (2019). Carrying out streamlined routine data analyses with reports for observational studies: introduction to a series of generic SAS ® macros. F1000research 7, 1955.

31. Miura, G.I., Roignant, J.Y., Wassef, M., and Treisman, J.E. (2008). Myopic acts in the endocytic pathway to enhance signaling by the Drosophila EGF receptor. Development 135, 1913–1922.

32. Palmer, R.H., Fessler, L.I., Edeen, P.T., Madigan, S.J., McKeown, M., and Hunter, T. (1999). DFak56 is a novel Drosophila melanogaster focal adhesion kinase. The Journal of biological chemistry 274, 35621–35629.

33. Martin-Bermudo, M.D., and Brown, N.H. (1999). Uncoupling integrin adhesion and signaling: the betaPS cytoplasmic domain is sufficient to regulate gene expression in the Drosophila embryo. Genes & Development 13, 729–739.

34. Cerami, E., Gao, J., Dogrusoz, U., Gross, B.E., Sumer, S.O., Aksoy, B.A., Jacobsen, A., Byrne, C.J., Heuer, M.L., Larsson, E., et al. (2012). The cBio Cancer Genomics Portal: An Open Platform for Exploring Multidimensional Cancer Genomics Data. Cancer discovery 2, 401–404.

35. Gao, J., Aksoy, B.A., Dogrusoz, U., Dresdner, G., Gross, B., Sumer, S.O., Sun, Y., Jacobsen, A., Sinha, R., Larsson, E., et al. (2013). Integrative Analysis of Complex Cancer Genomics and Clinical Profiles Using the cBioPortal. Sci Signal 6, pl1.

36. Ding, L., Getz, G., Wheeler, D.A., Mardis, E.R., McLellan, M.D., Cibulskis, K., Sougnez, C., Greulich, H., Muzny, D.M., Morgan, M.B., et al. (2008). Somatic mutations affect key pathways in lung adenocarcinoma. Nature 455, 1069–1075.

37. Imielinski, M., Berger, A.H., Hammerman, P.S., Hernandez, B., Pugh, T.J., Hodis, E., Cho, J., Suh, J., Capelletti, M., Sivachenko, A., et al. (2012). Mapping the Hallmarks of Lung Adenocarcinoma with Massively Parallel Sequencing. CELL 150, 1107–1120.

38. Campbell, J.D., Alexandrov, A., Kim, J., Wala, J., Berger, A.H., Pedamallu, C.S., Shukla, S.A., Guo, G., Brooks, A.N., Murray, B.A., et al. (2016). Distinct patterns of somatic genome alterations in lung adenocarcinomas and squamous cell carcinomas. Nature genetics 48, 607–616.

39. Jordan, E.J., Kim, H.R., Arcila, M.E., Barron, D., Chakravarty, D., Gao, J., Chang, M.T., Ni, A., Kundra, R., Jonsson, P., et al. (2017). Prospective Comprehensive Molecular Characterization of Lung Adenocarcinomas for Efficient Patient Matching to Approved and Emerging Therapies. Cancer discovery 7, 596–609.

40. Vavalà, T., Monica, V., Iacono, M.L., Mele, T., Busso, S., Righi, L., Papotti, M., Scagliotti, G.V., and Novello, S. (2017). Precision medicine in age-specific non-small-cell-lung-cancer patients: Integrating biomolecular results into clinical practice-A new approach to improve personalized translational research. Lung cancer (Amsterdam, Netherlands) 107, 84–90.

